# Aberrant MYCN expression drives oncogenic hijacking of EZH2 as a transcriptional activator in peripheral T cell lymphoma

**DOI:** 10.1101/2022.03.31.486583

**Authors:** Marlies Vanden Bempt, Koen Debackere, Sofie Demeyer, Quentin Van Thillo, Nienke Meeuws, Sarah Provost, Nicole Mentens, Kris Jacobs, Olga Gielen, David Nittner, Seishi Ogawa, Keisuke Kataoka, Carlos Graux, Thomas Tousseyn, Jan Cools, Daan Dierickx

## Abstract

Peripheral T cell lymphoma (PTCL) is a heterogeneous group of hematological cancers arising from the malignant transformation of mature T cells. In a cohort of 28 PTCL cases, we identified recurrent overexpression of MYCN, a member of the MYC family of oncogenic transcription factors. Approximately half of all PTCL cases was characterized by a MYC expression signature. Inducible expression of MYCN in lymphoid cells in a mouse model caused T cell lymphoma that recapitulated human PTCL with a MYC expression signature. Integration of mouse and human expression data identified EZH2 as a key downstream target of MYCN. Remarkably, EZH2 was found to be an essential co-factor for the transcriptional activation of the MYCN-driven gene expression program, which was independent of methyltransferase activity, but dependent on phosphorylation by CDK1. MYCN-driven T cell lymphoma was sensitive to EZH2 degradation or CDK1 inhibition, which displayed synergy with FDA-approved HDAC inhibitors.

**Key points:**

- Transcriptomic analysis of PTCL tumors reveals recurrent MYCN overexpression and the presence of a MYC signature in 50% of PTCL cases
- EZH2 is a transcriptional cofactor for the MYCN-driven gene expression program, which confers sensitivity to HDAC inhibition

## Introduction

Peripheral T cell lymphoma (PTCL) is a heterogeneous group of rare but aggressive hematological cancers arising from the malignant transformation of mature post-thymic T cells. Its heterogeneity is apparent from the most recent WHO classification describing at least 29 different subtypes. Except for ALK-positive anaplastic large cell lymphoma (ALCL), PTCL patients currently face an unfavorable prognosis: approximately 75% of all PTCL patients relapse after first-line chemotherapy treatment. Consequently, 5-year survival rates are as low as 10-30%, which underscores the urgent medical need for new therapeutic options^1–3^. Efforts to fully characterize the genetic and transcriptomic landscape of the different PTCL entities are currently ongoing. In line with this, we have previously performed whole transcriptome sequencing of a cohort of 28 clinical PTCL cases to identify novel oncogenic fusion genes^4^.

MYCN is a transcription factor of the MYC oncogene family, and its expression is typically low or absent in adult tissues^5^. While MYCN was previously found to be amplified and/or aberrantly overexpressed in several solid tumors, including neuroblastoma, prostate cancer and breast cancer^6^, it was not yet described as an oncogene in PTCL. The oncogenic role of MYCN has already been extensively studied in neuroblastoma, where amplification of MYCN is an initiating event present in 50% of high-risk patients^7^. Deregulated MYCN has been shown to invade tissue-specific promoter and enhancers, leading to the activation of oncogenic transcriptional programs^8^, and this MYCN-induced cell state was shown to be regulated by a transcriptional core regulatory circuitry consisting of a small number of transcription factors^9^. Consequently, MYCN-driven neuroblastoma was shown to be sensitive to BET inhibition^10^. Furthermore, a genome-wide CRISPR screen identified a specific dependency of MYCN-driven neuroblastoma on EZH2^11^. Indeed, EZH2 was found to directly interact with MYCN, thereby inducing its stabilization by protecting it from ubiquitination, and EZH2 depletion leads to downregulation of the MYCN-driven transcriptional program independent of its methyltransferase activity^12^.

The oncogenic function of EZH2 has been mostly associated with its canonical function as the enzymatic component of the PRC2 complex driving gene repression. Remarkably, in multiple cancer types, EZH2 was found to have a non-canonical function outside of the PRC2 complex as a transcriptional activator via its partially disordered transactivation domain that can be unlocked through cancer-specific phosphorylation^13,14^. This was found to be mostly independent of the methyltransferase activity, which implies that treatment strategies beyond EZH2 methyltransferase inhibitors, such as EZH2 degraders^15,16^, need to be considered for EZH2-overexpressing tumors.

In this study, we uncovered MYCN as an oncogenic driver in PTCL, and we identified EZH2 as an important cofactor for transcriptional activation of MYCN target genes.

## Methods

### Patient material

PTCL patient samples were collected retrospectively from the tumor banks of UZ Leuven and the CHU Mont-Godinne and prospectively in UZ Leuven. For prospectively obtained samples, we obtained informed consent from all patients. All cases were reviewed by two hematopathologists.

### Mice

We used 6- to 12-week old in-house bred male or female C57BL/6J mice, Tg(CD4-Cre) or Tg(CD2-iCre) mice as donors for the bone marrow transplant experiments, and female C57BL/6J mice as recipient mice for primary/secondary/tertiary transplants. Mouse experiments were approved and supervised by the KU Leuven ethical committee and conducted according to EU legislation (Directive 2010/63/EU).

Mice were housed in individually ventilated cages with a temperature between 18-23°C and humidity between 40-60% in SPF or semi-SPF conditions in the KU Leuven animal facility.

### Bone marrow transplant assays

For bone marrow transplant assays, hematopoietic stem/progenitor cells (HSPCs) were harvested from 6-12 weeks old in-house bred BL/6 mice (WT or transgenic CD2 Cre/CD4 Cre mice) using the EasySep Mouse Hematopoietic Progenitor Cell Isolation Kit (Stem Cell Technologies). HSPCs were transduced with retrovirus for expression of the desired oncogenes, and subsequently injected through tail-vein injection into recipient BL/6 mice that were irradiated 3-20h before injection (5 Gy).

Secondary and tertiary transplants were established using 6-12 weeks old female in-house bred BL/6 recipient mice. 500 000 – 1 000 000 spleen or thymus cells were injected through tail-vein injection into recipient mice that were irradiated 3-20h before injection (2.5 Gy). The survival of the mice was recorded daily.

### Flow cytometry

Single-cell suspensions were prepared from spleen, thymus, lymph nodes and bone marrow. The single cells were incubated in red blood cell lysis buffer (150 mM NH_4_Cl, 0.1 mM EDTA, 10 mM KHCO_3_) for 10 minutes prior to staining. Cells were then washed with PBS and stained in the dark for 20 minutes at 4°C. Next, stained cells were washed with PBS and then analyzed on a MACSQuant Vyb (Miltenyi).

Annexin V stainings were performed using the APC Annexin V Apoptosis Detection Kit (BioLegend) according to the manufacturer’s instructions.

Intracellular stainings were performed using the Foxp3 / Transcription Factor Staining kit (Thermo Fischer Scientific) according to the manufacturer’s instructions.

Data were analyzed with the FlowJo software (Tree Star). Antibodies are listed in Table S1.

### ChlPmentation ChIP-sequencing

ChlPmentation ChlP-seq was performed as described previously^17–19^. 20-50 million primary spleen cells were washed with PBS and cross-linked with 1% formaldehyde for 10 min and then quenched by addition of glycine. For nuclei isolation, cells were resuspended in 1X RSB buffer (10 mM Tris pH7.4,10 mM NaCl, 3 mM MgCl2) and left on ice for 10 min. Cells were collected and resuspended in RSBG40 buffer (10 mM Tris pH7.4, 10 mM NaCl, 3 mM MgCl2, 10% glycerol, 0.5% NP40) with 1/10 v/v of 10% detergent (3.3% w/v sodium deoxycholate, 6.6% v/v Tween-40). Nuclei were collected and resuspended in L3B buffer (10 mM Tris-Cl pH 8.0, 100 mM NaCl, 1 mM EDTA, 0.5 mM EGTA, 0.1% Na-Deoxycholate, 0.5% N-Lauroylsarcosine, 0.2% SDS). Chromatin was fragmented to 200-400 bp using a Bioruptor (Diagenode) for 20-25 cycles (30 s on, 30 s off, High). The chromatin was supplemented with 1% Triton-X100 after fragmentation. Antibodies (listed in Table S4) were pre-conjugated to magnetic protein A/G beads (Millipore). Chromatin immunoprecipitation was carried out overnight. Tagmentation and library preparation was performed using the Nextera DNA library prep kit (Illumina). DNA was purified using triple sided SPRI bead clean-up (Agencourt AMPure Beads, Beckman Coulter).

### Dual luciferase assay

HEK293T cells at 80-90% confluency in a 12-well plate were transfected using 6 μL GeneJuice Transfection Reagent (Sigma-Aldrich, 70967) and 100 μL RPMI 1640 medium (ThermoFisher) per well. Each well was transfected with 500 ng of pGL4.27 (Promega), in which we cloned a fragment of the murine *Aurka* promoter **(Sup. Fig. 3D)** or no response element, 25 ng of renilla luciferase control reporter vector (RL-TK, Promega) and the respective plasmids for the expression of each transcription factor (MYCN, EZH2 and variants) or empty vector in triplicates. Cells were lysed after 24 hours with Passive Lysis Buffer and the luciferase signal was measured on a Victor multilabel plate reader (PerkinElmer) using the Dual-Luciferase Reporter Assay System (Promega). The signal was normalized to the Renilla luciferase activity.

### *Ex vivo* inhibitor treatments

Cells harvested from the spleen of mice that developed MYCN-driven TCL were seeded into 96-well plates in RPMI 1640 supplemented with 20% fetal calf serum, 2 mM L-glutamine, non-essential amino acids solution (Gibco), 50 μM β-mercapto-ethanol, primocin (InVivoGen), 100 ng/mL interleukin-2 and 50 ng/mL interleukin-7. The compounds were added in a randomized fashion using a D300e digital dispenser (Tecan) and the DMSO concentration was normalized. All compounds are listed in Table S6. Plates were incubated overnight and cell proliferation was measured after 24h using the ATPlite luminescence system (PerkinElmer) using a Victor multilabel plate reader. For combination treatments with multiple compounds, data were analyzed using the SynergyFinder software^20^.

### Data analysis

All statistical analyses were performed using Prism software (Graphpad software, CA, USA). For analysis of the mouse data, survival was calculated using the Kaplan-Meier method.

For qRT-PCR analyses, data are expressed as the mean ± standard deviation (SD). Comparisons between two groups were performed using one-way ANOVA with corrections for multiple comparisons.

Paired-end RNA sequencing data was collected from the patient samples, while all other RNA sequencing data is single 3’ end transcriptome data. Libraries were sequenced on a HiSeq 4000 with 125bp single-end reads (Illumina). All RNA-seq data was first cleaned with fastq-mcf from ea-utils, after which a quality control was performed with FastQC. Subsequently the reads were mapped to the respective reference genomes, i.e. either GRCh38/hg38 or GRCm38/mm10, with the HISAT2 software. Further processing of the reads was executed with the SAMtools package and the number of reads per transcript was determined with HTSeq-count. The R-package DESeq2 was used to determine the significantly differentially expressed genes. Heatmaps and expression plots were constructed with R. Gene set enrichment analyses were performed with the GSEA software from the Broad Institute. Possible regulatory transcription factor motifs were determined with i-CisTarget^21^.

ChIPmentation sequencing data with either specific antibodies against transcription factor or histone marks, was preprocessed similarly as the RNA-seq data, i.e. cleaning with fastq-mcf and quality control with FastQC. Subsequently, these cleaned read were mapped to the respective reference genome (GRCm38/mm10) with the Bowtie2 software, after which the reads were further processed with SAMtools. The ChIP signals were normalized with the deepTools package (bamCoverage) and the peaks were called with MACS2. These peaks were then annotated with a custom script. The centered heatmaps were constructed with the deepTools package (computeMatrix, plotHeatmap).

The RNA-sequencing and ChIP-sequencing data were deposited in the Gene Expression Omnibus (GEO) database with accession number GSE198126.

## Results

### MYCN is aberrantly expressed in PTCL and drives a MYC target gene expression signature

To gain a deeper understanding of the mechanisms that drive PTCL development, we carried out RNA-sequencing (RNA-seq) on 28 PTCL samples, including 23 PTCL, not otherwise specified (PTCL-NOS) cases, 2 ALCL cases and 3 Follicular T cell lymphoma (FTCL) cases^4^. In 5 of these 28 PTCL cases (18%), we identified aberrant overexpression of *MYCN* **(Fig. 1A).** In addition, we were able to identify additional cases with *MYCN* overexpression in publicly available PTCL datasets^22–24^ **(Fig. 1B),** while *MYCN* overexpression was extremely rare in diffuse large B cell lymphoma (DLBCL) **(Fig. S1A).**

**Figure 1:**
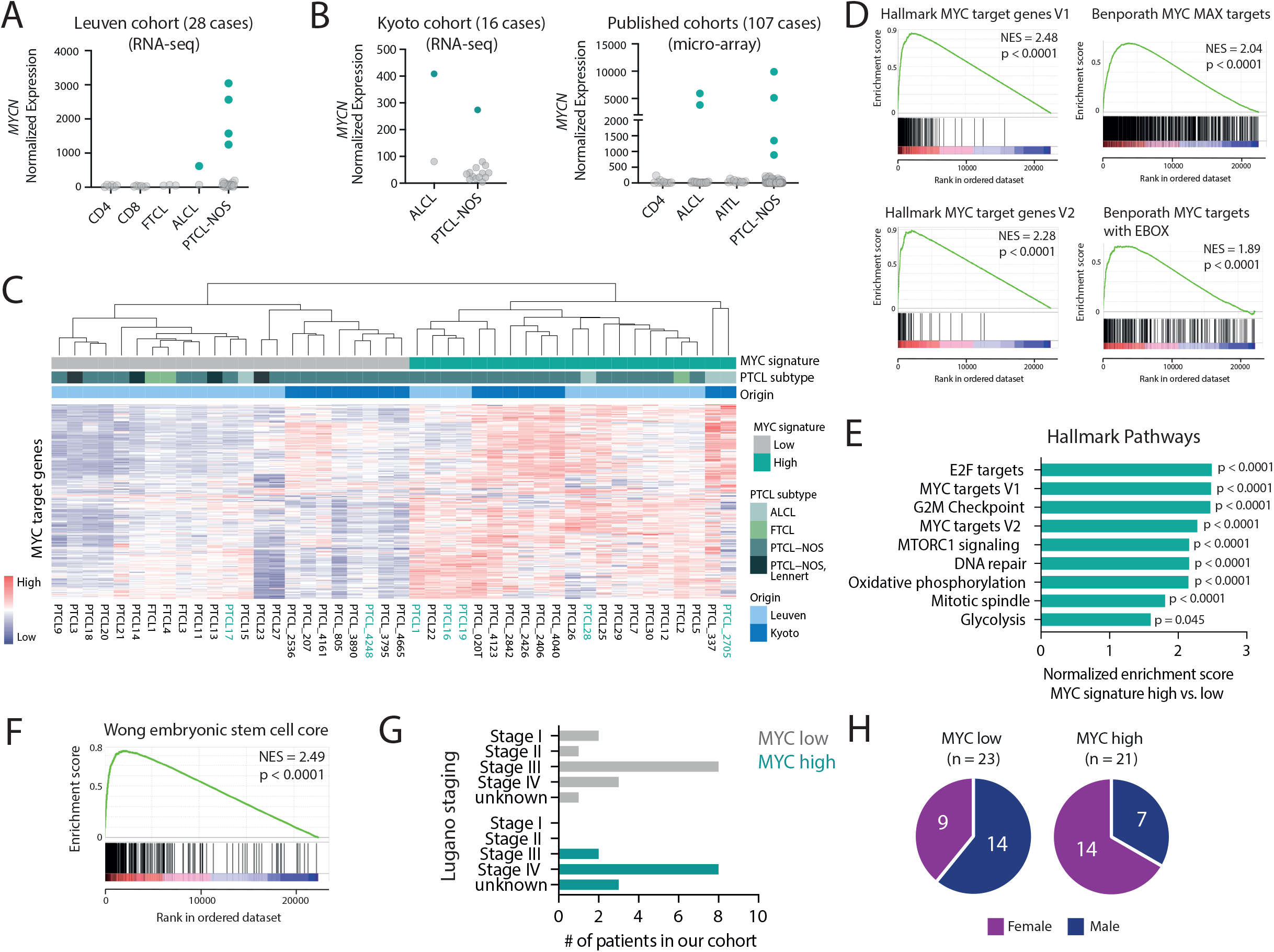
MYCN is aberrantly overexpressed in PTCL. (A) Normalized counts for MYCN in the Leuven PTCL cohort of 28 cases. (B) Normalized counts for MYCN in three published datasets (RNA-seq ^24^ and micro-array ^22,23)^. (C) Heatmap showing expression of MYC target genes from the Hallmark gene sets in PTCL cases from the Leuven and Kyoto cohorts. Samples indicated in green harbor overexpression of MYCN. (D) Gene set enrichment analysis (GSEA) showing enrichment of different MYC target gene sets in the differentially expressed genes of PTCL with a high MYC signature compared to PTCL with a low MYC signature. NES = normalized enrichment score. (E) Normalized enrichment scores for different Hallmark gene sets in MYC high versus low PTCL. (F) GSEA showing enrichment of an embryonic stem-cell like signature as described in Wong et al., 2008. (G) Lugano staging for the PTCL patients in the Leuven cohort. (H) Pie chart representing the sex of the PTCL patients in the MYC low (left) and MYC high (right) subgroups. The difference between the female presence in the two groups was calculated using a hypergeometrical distribution (p = 0,016). See also Figure S1.

We next investigated the expression of known MYC target genes in PTCL. Unsupervised clustering showed that approximately half of the PTCL cases are characterized by a high MYC signature **(Fig. 1C, D),** and these cases showed the highest expression levels of *MYCN, MYC, MYB, MYBL1* and/or *MYBL2* **(Fig. S1B).** Remarkably, the presence of a MYC signature was mutually exclusive with a diagnosis of Lennert lymphoma, a lymphoepithelioid variant of PTCL-NOS **(Fig. 1C, Fig. S1C).** PTCL cases with a high MYC signature showed strong positive enrichment of gene sets related to E2F targets, cell cycle checkpoints and DNA repair **(Fig. 1E, Fig. S1D),** indicative of a proliferative state. We also observed strong enrichment of a gene set representing an embryonic stem cell-like state that was previously associated with MYC-induced cancers^25^ **(Fig. 1F).** Our MYC signature classification did not replicate the classification based on *TBX21* or *GATA3* expression^22^ **(Sup. Fig. 1E,F).** Clinically, cases with a high MYC signature presented with more advanced disease, as reflected by the Lugano staging at diagnosis **(Fig. 1G)** and were more often female (p = 0,016, **Fig. 1H).** No difference in age at diagnosis was observed **(Fig. S1G).**

Taken together, we identified *MYCN* as a novel recurrent oncogene in PTCL and uncovered the presence of a high MYC signature in approximately half of the PTCL cases, which correlates with a more aggressive and proliferative disease.

### MYCN overexpression drives the development of T cell lymphomas in a bone marrow transplant mouse model

Next, we aimed to further explore the oncogenic potential of *MYCN* overexpression *in vivo. We* used a bone marrow transplant assay to generate mice expressing *MYCN* in a subset of hematopoietic stem/progenitor cells (HSPCs). However, constitutive expression of *MYCN* in HSPCs led to a rapid development of myeloid leukemia, as previously described ^26^ **(Fig. S2A, B)**. To model T cell lymphoma, we restricted *MYCN* expression to the lymphoid lineage by using a Cre-inducible retroviral vector^18^ in combination with CD2 Cre or CD4 Cre transgenic mice as donors **(Fig. 2A).** Recipient mice transplanted with CD2 Cre HSPCs transduced with the inducible *MYCN* vector (also expressing GFP) developed lymphomas with a B or T cell phenotype with a median survival of 56 days **(Fig. 2B, Fig. S2C-E).** Both T cell and B cell lymphomas were transplantable into secondary recipients with a short latency **(Fig. 2C).** Recipient mice transplanted with CD4 Cre HSPCs transduced with the inducible *MYCN* vector uniquely developed T cell lymphomas with a longer median latency (202 days) **(Fig. 2D, Fig. S2G).**

**Figure 2:**
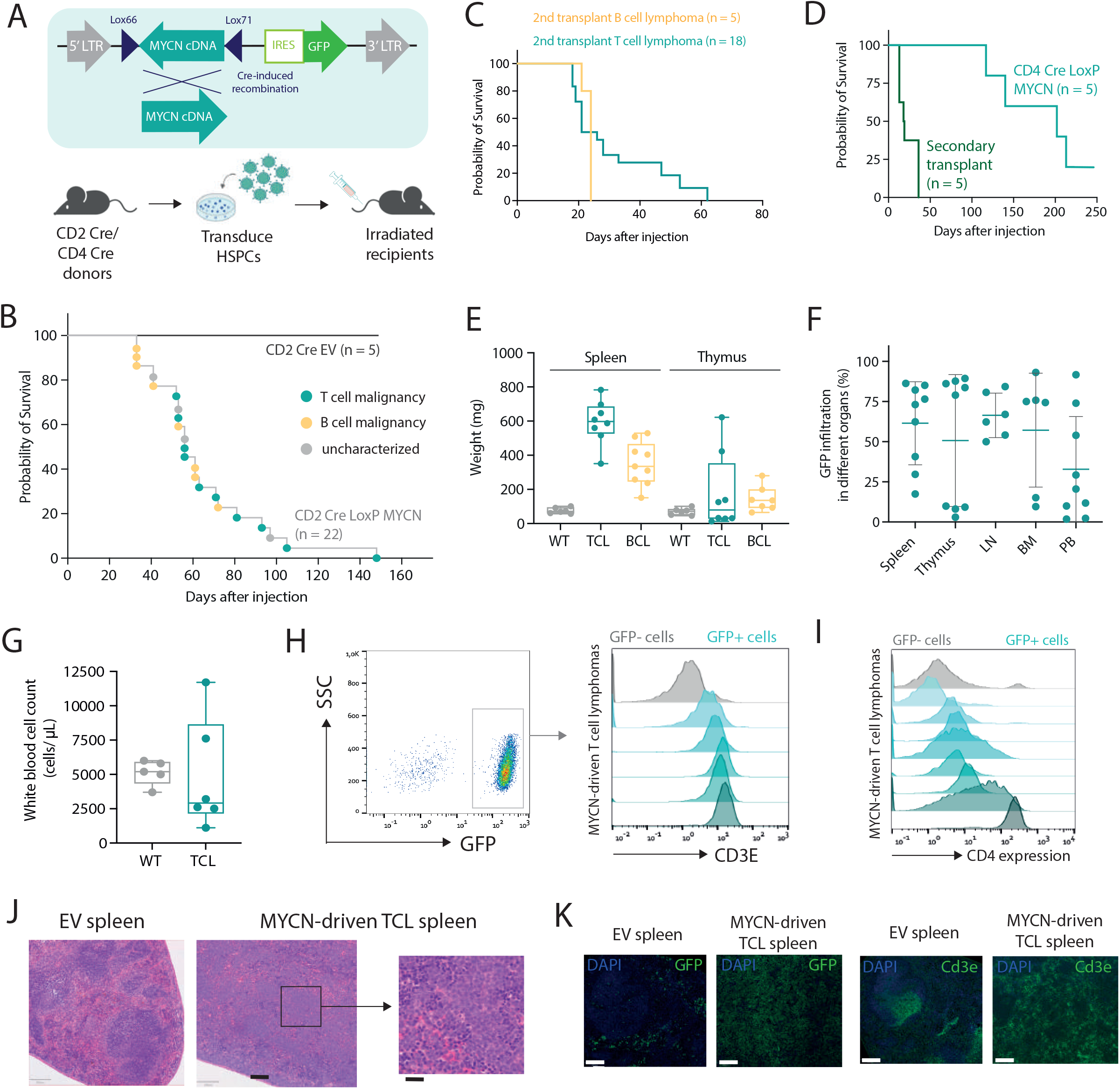
MYCN drives PTCL in a bone marrow transplant model. (A) Schematic overview of the bone marrow transplant assay strategy used to restrict MYCN expression to the lymphoid lineage. (B) Kaplan-Meier overall survival curve for mice transplanted with CD2 Cre HSPCs transduced with empty vector (n = 5) or the LoxP MYCN retroviral vector (n = 22). (C) Kaplan-Meier overall survival curve for secondary transplants of MYCN-driven B cell lymphoma (n = 5) or MYCN-driven T cell lymphoma (n = 18). (D) Kaplan-Meier overall survival curve for mice transplanted with CD4 Cre HSPCs transduced with the LoxP MYCN retroviral vector (n = 5), and secondary transplants from the resulting T cell lymphomas (n = 5). (E) Spleen and thymus weights in mice that develop MYCN-driven B cell or T cell lymphomas or WT mice. (F) Infiltration of GFP-positive cells in different organs from mice that develop MYCN-driven TCL. (G) White blood cell counts of WT mice or recipient mice transplanted with CD2 Cre cells transduced with the LoxP MYCN retroviral vector at end stage disease (T cell lymphoma). (H) Representative FACS analysis showing CD3e expression in GFP-positive spleen cells of mice that developed MYCN-driven TCL. (I) Representative FACS analysis showing CD4 expression in GFP-positive spleen cells of mice that developed MYCN-driven TCL. (J) H&E stainings on a spleen from a mouse transplanted with HSPCs expressing empty vector (GFP) and a spleen from the murine MYCN-driven TCL. Scale bars indicate 200 μm (black) or 20 μm (white). (K) Immunohistochemistry staining showing GFP (top) or Cd3e (bottom) in a spleen from a mouse transplanted with HSPCs expressing empty vector (EV) and a spleen from a mouse that developed MYCN-driven TCL. Scale bars indicate 200 μm. See also Figure S2.

The mice that developed T cell lymphomas presented with splenomegaly **(Fig. 2E),** and some animals developed paraplegia. We typically observed high infiltration of GFP-positive cells in spleen and lymph nodes, and more variable levels of infiltration in thymus, bone marrow and peripheral blood **(Fig. 2F),** but no strong increase in white blood cell count **(Fig. 2G).** Phenotypic analysis revealed that the malignant T cells were expressing surface CD3E **(Fig. 2H)** and showed variable levels of CD4 expression **(Fig. 2I),** but low or absent CD8 expression **(Fig. S2I).** Histopathological analysis showed a profound loss of spleen architecture and infiltration of mitotic cells and confirmed high levels of CD3E and GFP expression in the affected organs **(Fig. 2J, K).**

To further investigate the transcriptional programs underlying MYCN-driven T cell lymphoma (TCL), we performed a global gene expression analysis using RNA-seq on six murine MYCN-driven TCL samples and on CD4 T cells isolated from the spleens of three WT mice **(Fig. S3A).** Motif analysis revealed a strong enrichment of E2F and MYC motifs in genes that are upregulated in murine MYCN-driven TCL **(Fig. 3A).** In line with this, GSEA revealed strong enrichment of gene sets containing E2F target genes, MYC target genes, gene sets related to cell cycle progression, DNA repair, and MYC-driven sternness^25^ **(Fig. 3B, C),** as was the case in the human PTCL cases characterized by a high MYC signature. Malignant cells expressed high levels of *Pdcd1*, but low levels of the T_FH_ markers *Icos, Cxcr5* and *Bcl6* and the ALCL markers *Batf3* and *Irf4* **(Fig S3B),** indicating a PTCL-NOS phenotype. Expression analysis of T cell receptor variable genes confirmed the oligoclonal nature of the lymphomas **(Fig. S3C).**

**Figure 3:**
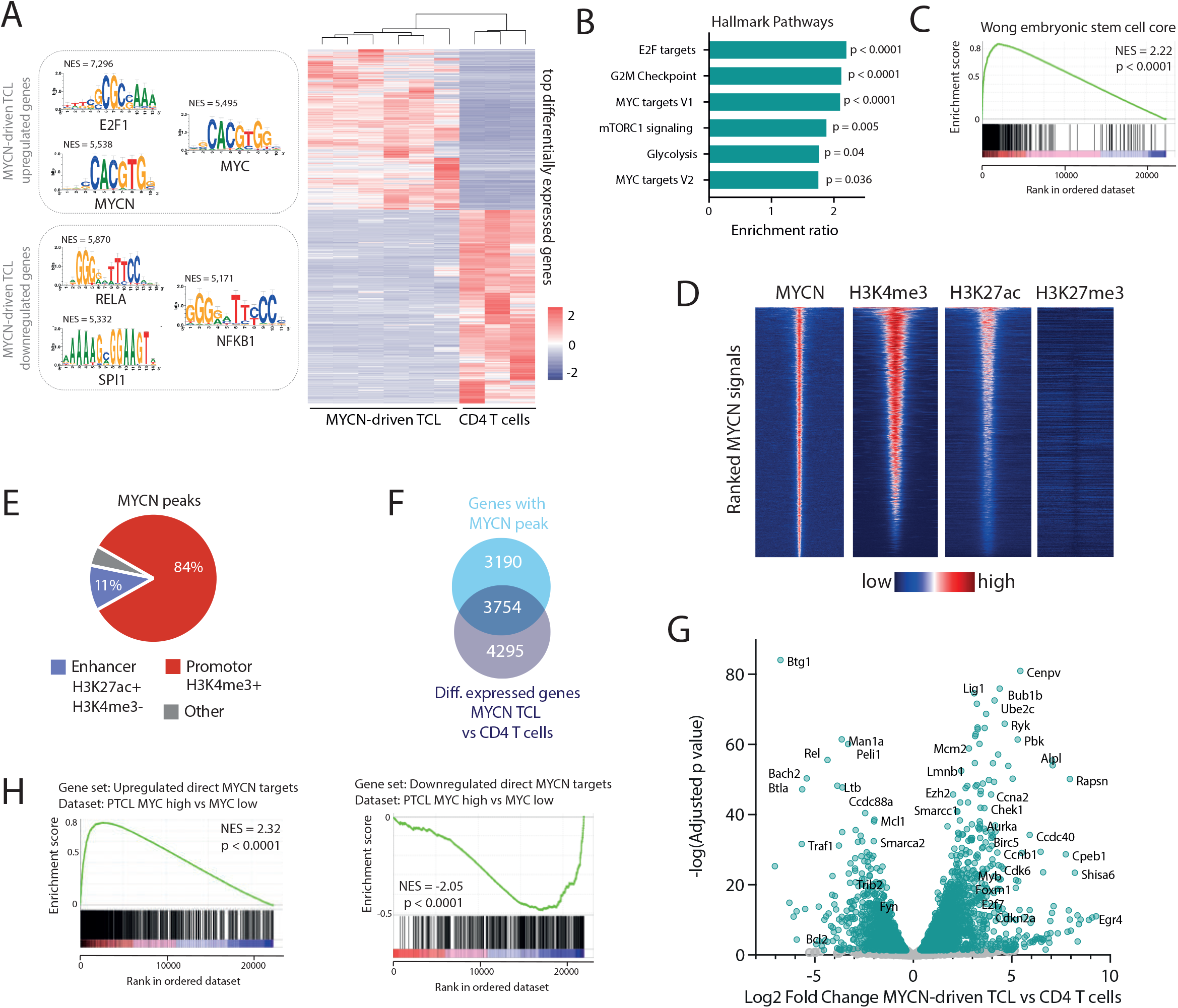
Murine MYCN-driven T cell lymphoma recapitulates human PTCL with a high MYC signature. (A) Heatmap showing the top differentially expressed genes in MYCN-driven TCL compared to normal CD4 T cells and depicting the transcription factor binding motifs of the top transcription factors identified by i-CisTarget in the up- and downregulated genes. NES = normalized enrichment score. (B) Normalized enrichment scores for different Hallmark gene sets in MYCN-driven TCL compared to normal CD4 T cells. (C) GSEA showing enrichment of an embryonic stem-cell like signature as described in Wong et al., 2008. (D) Centered read-density heatmaps showing binding locations of MYCN, H3K4me3, H3K27ac and H3K27me3 in MYCN-driven TCL. Heatmaps centered and ranked on MYCN signal strength. (E) ChIP-seq peak occurrence of MYCN in relation to promoter regions (H3K4me3+) and enhancer regions (H3k27ac+/ H3K4me3-). (F) Venn diagram showing the total amount of ChIP-seq peaks of MYCN in promotors of genes that are differentially expressed in MYCN-driven TCL compared to normal CD4 T cells. (G) Volcano plot showing the Log2 fold change of direct MYCN target genes in MYCN-driven TCL compared to CD4 T cells. (H) GSEA to show negative enrichment of downregulated direct MYCN target genes (FC < - 1.5) and positive enrichment of upregulated direct MYCN target genes (FC > 1.5) in the differentially expressed genes in PTCL with a high versus low MYC signature. See also Figure S3.

Subsequently, we performed chromatin immunoprecipitation followed by sequencing (ChIP-seq) for MYCN and various histone marks. MYCN was found to bind most often in promotor regions, and MYCN binding locations were enriched in H3K4me3 and H3K27ac **(Fig. 3D, E).** We integrated our ChIP-seq and RNA-seq data to determine which genes are directly regulated by MYCN (e.g. genes that are bound by MYCN in their promoter region and that are differentially expressed in MYCN-driven TCL compared to non-malignant CD4 T cells). We identified 3754 direct MYCN target genes, from which 2429 genes were upregulated and 1325 genes were downregulated **(Fig. 3F, G).** These direct MYCN target genes were also found to be enriched in differentially expressed genes in PTCL with a high versus a low MYC signature **(Fig. 3H).**

Altogether, these results show that MYCN acts as a strong oncogene in the lymphoid lineage, and that this mouse model of MYCN-driven TCL reflects human PTCL with a high MYC signature.

### Ezh2 is implicated in transcriptional activation of direct MYCN target genes

Within the most significantly upregulated direct MYCN target genes that were also differentially expressed in human PTCL with a high MYC signature, we identified genes related to cell cycle and DNA replication **(Fig. 4A).** Surprisingly, *Ezh2*, but not *Suz12, Eed* or *Ezh1*, was found amongst the most significantly upregulated direct MYCN target genes **(Fig. 4B-D, Fig. S4A, B).** Similarly, we observed increased *EZH2* expression levels in human PTCL with a high MYC signature **(Fig. 4E).** Interestingly, Ezh2 peaks were found both in repressed regions (defined by H3K27me3 marks) where co-binding with Suz12 was observed, as well as in active promotor regions (defined by H3K4me3 and H3K27ac marks, p300 binding and active gene expression). Co-occupancy of Ezh2 with MYCN and p300 in active promotor regions was not accompanied by Suz12 or the repressive H3K27me3 mark, indicating that Ezh2 binds these regions independent of the PRC2 complex **(Fig. 4F).** Remarkably and unexpectedly, when looking at direct MYCN target genes, Ezh2 binding was mostly associated with H3K27ac rather than H3K27me3 **(Fig. S4C)** and these activating Ezh2 peaks were enriched for the same gene sets as MYCN-driven TCL and for a gene set containing EZH2 targets that preserve stem cell potential in hematopoietic stem cells ^27^ **(Fig. 4G, H).** Together, these findings suggest that Ezh2 acts as a transcriptional cofactor for MYCN.

**Figure 4:**
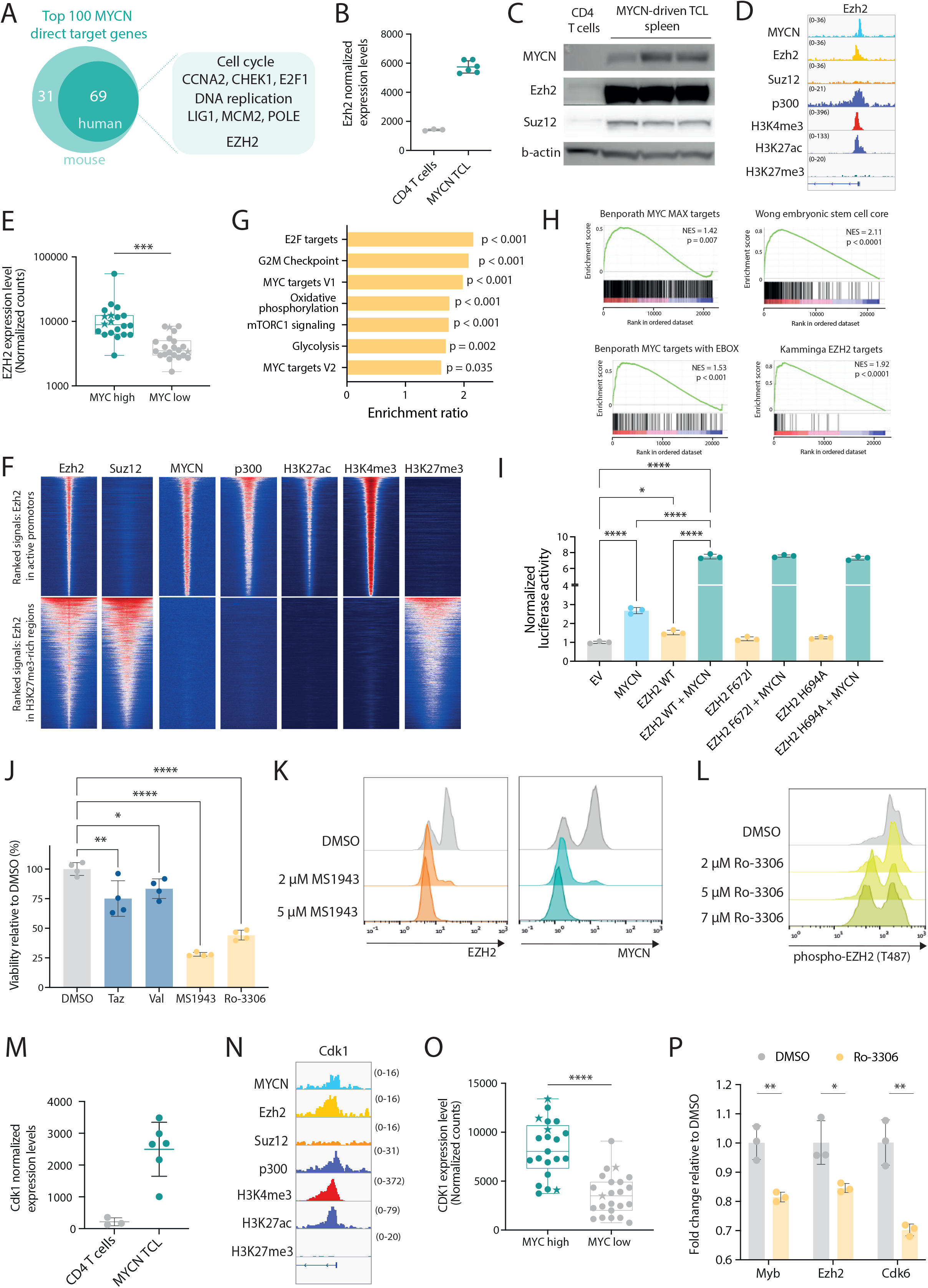
EZH2 is a co-factor for transcriptional activation together with MYCN. (A) Schematic representation of the top 100 MYCN direct target genes. (B) Normalized expression levels *of Ezh2* in CD4 T cells and in MYCN-driven lymphoma cells. (C) Western blot showing expression levels of MYCN, Ezh2 and Suz12 in normal CD4 T cells and in MYCN-driven TCL spleen cells. (D) Representative ChIP-seq tracks for the *Ezh2* promoter showing binding of MYCN, Ezh2, Suz12, p300 and histone marks H3K4me3, H3K27ac and H3K27me3 in MYCN-driven TCL. (E) Normalized counts for *EZH2* from the PTCL-NOS and FTCL cases from the Leuven cohort and the Kyoto cohort. Patients with overexpression of MYCN are indicated with stars. (F) Centered read-density heatmaps showing binding locations of Ezh2, Suz12, MYCN, p300, H3K27ac, H3K4me3 and H3K27me3 in MYCN-driven TCL. Heatmaps centered and ranked on Ezh2 signal strength in active promoters (top) or in regions enriched for H3K27me3 (bottom). (G) Normalized enrichment scores for different Hallmark gene sets in the differentially expressed direct EZH2 + MYCN target genes. (H) GSEA showing enrichment of different MYC target gene sets and sternness-associated gene sets in the differentially expressed direct EZH2+MYCN target genes. (I) Luciferase assay in HEK293T cells showing transcriptional activation of Firefly luciferase by MYCN and/or EZH2 (wild type or methyltransferase-inactivating mutants). (J) 24h *ex vivo* treatment of MYCN-driven TCL spleen cells with 2 μM of the EZH2 methyltransferase inhibitors Tazemetostat (Taz) and Valemetostat (Val), the EZH2 degrader MS1943, or the CDK1 inhibitor Ro-3306. (K) FACS analysis of EZH2 and MYCN in MYCN-driven TCL spleen cells treated *ex vivo* with DMSO or MS1943 for 12h. (L) FACS analysis of phospho-EZH2 (T487) in MYCN-driven TCL spleen cells treated *ex vivo* with DMSO or Ro-3306 (CDK1 inhibitor) for 8h. (M) Normalized expression levels of *Cdk1* in CD4 T cells and in MYCN-driven lymphoma cells. (N) Representative ChIP-seq tracks for the *Cdk1* promoter showing binding of MYCN, Ezh2, Suz12, p300 and histone marks H3K4me3, H3K27ac and H3K27me3 in MYCN-driven TCL. (O) Normalized counts for *CDK1* from the PTCL cases from the Leuven cohort and the Kyoto cohort. (P) qRT-PCR analysis of MYCN/EZH2 target gene expression in MYCN-driven TCL cells treated *ex vivo* for 8h with 4 μM Ro-3306. Data are represented as mean± SD. See also Figure S4.

To further investigate the role of EZH2 in MYCN transcriptional activity, we performed a luciferase reporter experiment with a luciferase expression construct under control of the *Aurka* promoter, a region that we identified as strongly bound by MYCN and EZH2 **(Fig. S4D).** Expression of MYCN alone caused increased luciferase activity, which was further enhanced when MYCN was co-expressed with EZH2. Mutations in EZH2 that inactivate the methyltransferase activity had no effect on the transcriptional activity in the luciferase reporter assay, indicating that the enzymatic activity of EZH2 is not required **(Fig. 4I, Fig. S4E, F).** In line with this, MYCN-driven TCL cells showed only a slight sensitivity to inhibitors that block the methyltransferase activity of Ezh2, yet they were strongly sensitive to targeted Ezh2 degradation by MS1943 **(Fig. 4J, Fig. S4G, H).** Furthermore, Ezh2 depletion induced MYCN destabilization and subsequent degradation **(Fig. 4K),** as previously reported in neuroblastoma^12^.

Subsequently, we performed immunoprecipitation of Ezh2 in MYCN-driven TCL cells, followed by mass spectrometry. We uncovered phosphorylation of Ezh2 at Threonine 487 **(Fig. 4L, Fig. S4I),** a post-translational modification catalyzed by CDK1, which was previously associated with decreased interaction with PRC2 complex members SUZ12 and EED ^28^. Interestingly, *Cdk1* was found to be a direct target gene of MYCN and Ezh2 in MYCN-driven TCL **(Fig. 4M, N).** Moreover, *CDK1* was significantly upregulated in human PTCL with a high MYC signature **(Fig. 4O).** In line with this, treatment with the CDK1 inhibitor Ro-3306 reduced phosphorylation of EZH2 **(Fig. 4L)** and diminished the expression of genes directly regulated by MYCN and EZH2 **(Fig. 4P).** Moreover, MYCN-driven TCL cells were sensitive to CDK1 inhibition **(Fig. 4J).**

Collectively, these data show that, in the context of PTCL, EZH2 plays an important role outside of the PRC2 complex as a transcriptional co-activator together with MYCN, thereby reinforcing the transcriptional programs established by MYCN overexpression.

### MYCN-driven T cell lymphoma is addicted to high levels of EZH2 and displays a strong sensitivity for HDAC inhibition

Having established EZH2 as a strong transcriptional co-factor for MYCN, we wanted to assess EZH2 inhibition as therapeutic approach for MYCN-driven TCL. Therefore, we performed an *ex vivo* treatment of MYCN-driven TCL cells with the EZH2 selective degrader MS1943, and we could observe a strong dependency on Ezh2 **(Fig. 5A).** Similarly, we inhibited Ezh2 phosphorylation on T487 using the CDK1 inhibitor Ro-3306, and this induced apoptosis in MYCN-driven TCL cells **(Fig. 5B, C).** As expected, MYCN-driven TCL cells also showed sensitivity to BET inhibition **(Fig. 5D).**

As EZH2 degraders are not yet ready to use in clinical practice, we subsequently investigated the potential of clinically available inhibitors. The class I HDAC inhibitors romidepsin and belinostat are already FDA-approved for treatment of PTCL, but only induce durable response in a subset of patients and no reliable biomarkers are available yet^1^. We observed upregulation of class I HDACs in PTCL cases with a high MYC signature **(Fig. 5E),** and in line with this, Hdac1 and Hdac2 were found to be direct MYCN targets in MYCN-driven TCL **(Fig. 5F, G).** Therefore, we investigated the sensitivity of MYCN-driven TCL to HDAC inhibition. The class I HDAC inhibitors romidepsin and belinostat both induced strong inhibitory effects at low concentrations in MYCN-driven TCL cells **(Fig. 5H, Fig. S5A),** which resulted in downregulation of MYCN/EZH2 target genes **(Fig. 5I),** and induction of apoptosis after only 8h of treatment **(Fig. 5C).**

**Figure 5:**
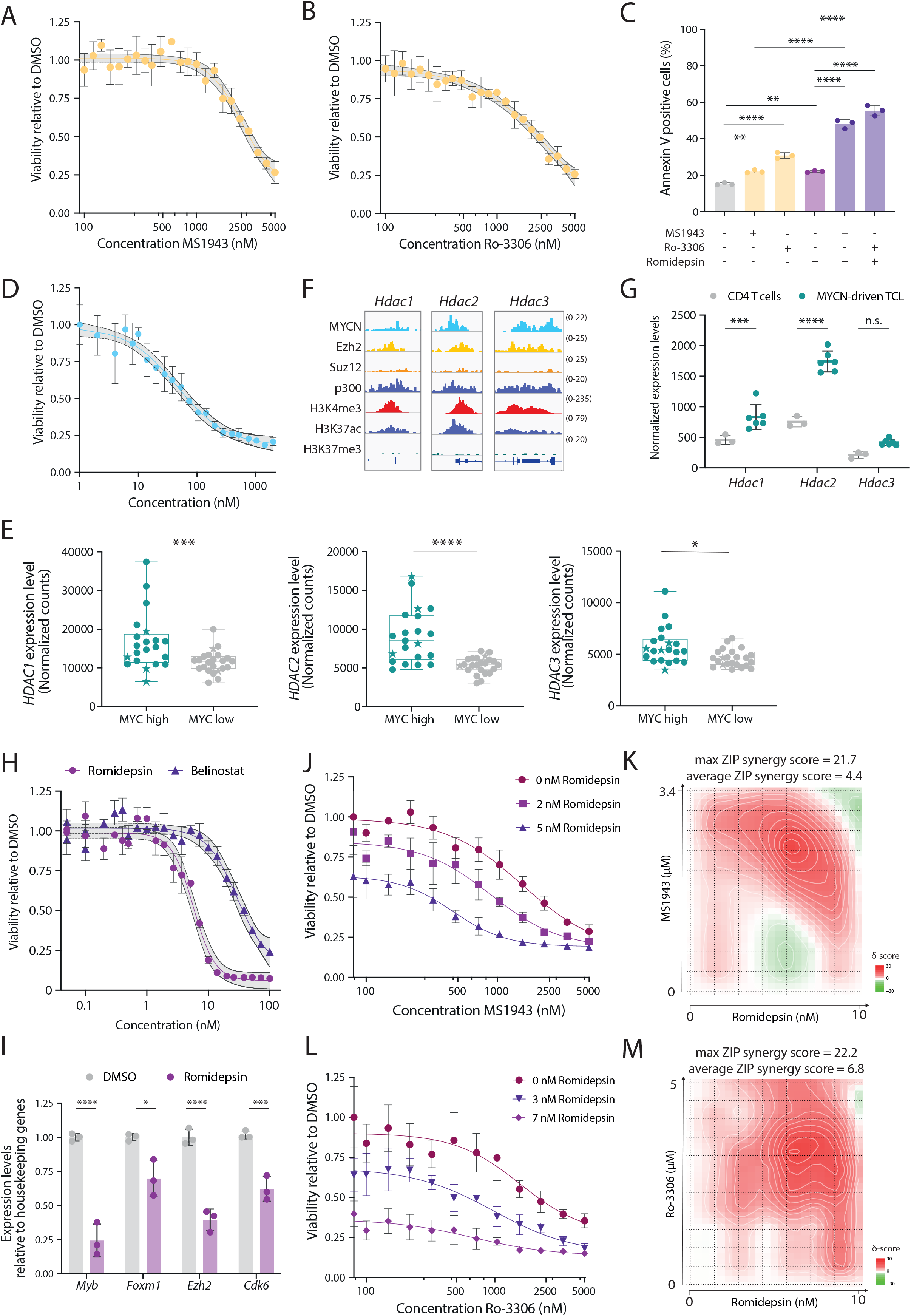
MYCN-driven TCL is sensitive to EZH2 depletion and HDAC inhibition. (A) Dose response curve for 24h treatment with the EZH2 degrader MS1943 on *ex vivo* cultured MYCN-driven TCL cells. (B) Dose response curve for 24h treatment with the CDK1 inhibitor Ro-3306 on *ex vivo* cultured MYCN-driven TCL cells. (C) Annexin V staining of MYCN-driven TCL cells treated *ex vivo* for 8h with DMSO, 10 nM romidepsin, 5 μM Ro-3306, 5 μM MS1943, or a combination of these inhibitors. (D) Dose response curve for 24h treatment with the BET inhibitor JQ1 on *ex vivo* cultured MYCN-driven TCL cells. (E) Normalized counts for *HDAC1, HDAC2* and *HDAC3* from the PTCL cases from the Leuven cohort and the Kyoto cohort. (F) Representative ChIP-seq tracks for the *Hdac1, Hdac2* and *Hdac3* promoter regions showing binding of MYCN, Ezh2, Suz12, p300 and histone marks H3K4me3, H3K27ac and H3K27me3 in MYCN-driven TCL. (G) Normalized expression levels of *Hdac1, Hdac2* and *Hdac3* in CD4 T cells and in MYCN-driven lymphoma cells. (H) Dose response curve for 24h treatment with the HDAC inhibitors romidepsin and belinostat on *ex vivo* cultured MYCN-driven TCL cells. (I) qRT-PCR analysis of MYCN/EZH2 target gene expression in MYCN-driven TCL cells treated *ex vivo* for 4h with 500 nM Romidepsin. Data are represented as mean± SD. (J) Dose response curve for 24h treatment with the EZH2 degrader MS1943 with or without simultaneous romidepsin treatment on *ex vivo* cultured MYCN-driven TCL cells. (K) Synergy matrix plot showing δ-scores for *ex vivo* cultured MYCN-driven TCL cells treated with romidepsin + MS1943 (max ZIP synergy score = maximal score for a specific dose combination; average ZIP synergy score = the average δ-score for the whole range of concentrations shown in the synergy matrix). (L) Dose response curve for 24h treatment with the CDK1 inhibitor Ro-3306 with or without simultaneous romidepsin treatment on *ex vivo* cultured MYCN-driven TCL cells. (M) Synergy matrix plot showing δ-scores for *ex vivo* cultured MYCN-driven TCL cells treated with romidepsin + Ro-3306 (max ZIP synergy score = maximal score for a specific dose combination; average ZIP synergy score = the average δ-score for the whole range of concentrations shown in the synergy matrix). See also Figure S5.

Finally, we combined HDAC inhibition with EZH2 degradation, CDK1 inhibition or BET inhibition. We calculated synergy using the zero interaction potential (ZIP) method ^20^, and observed synergistic effects between EZH2 degradation/dephoshorylation and HDAC inhibition **(Fig. 5J-M, Fig. S5B-E).**

## Discussion

PTCL is a heterogeneous and poorly characterized malignancy, for which very few treatment options other than chemotherapy are currently available. Despite recent advances in our understanding of the genetic and transcriptomic landscape of PTCL, no novel targeted treatment strategies have been developed, as it became apparent that PTCL tumors are characterized by many different aberrations that occur at a low frequency. For many of these genetic or transcriptomic alterations, their exact contribution to PTCL development is unknown.

In this study, we uncovered *MYCN* as a novel recurrent oncogenic driver in 18% of PTCL cases. The oncogenic role of MYCN was already extensively studied in other tumor types, such as neuroblastoma and neuro-endocrine prostate cancer, but its implication in PTCL was not previously described. Similar to neuroblastoma mouse models, where *MYCN* overexpression is restricted to noradrenergic neurons^29^, overexpression of *MYCN* in the lymphoid lineage was sufficient to drive development of both B and T cell malignancies. Although *MYCN* overexpression is activated during early lymphoid development in our bone marrow transplant models, recipient mice only develop mature T or B cell malignancies. This suggests that, while *MYCN* has been described as an oncogenic driver in T cell acute lymphoblastic leukemia^30^, additional factors are needed to block T cell differentiation, and that *MYCN* preferentially transforms mature lymphoid cells.

Intriguingly, we identified EZH2 as an essential co-factor for MYCN-driven transcriptional activation in T cell lymphomas. The dependency of MYCN-overexpressing tumors on EZH2 already became apparent from a genome-wide CRISPR screen in MYCN-amplified neuroblastoma, which revealed a specific dependency on PRC2 complex members EZH2, EED and SUZ12^11^. Moreover, MYCN was shown to directly activate EZH2 expression. Nevertheless, EZH2 binding in these tumors was strongly correlated with repressive histone marks, and the methyltransferase activity of EZH2 was found to be essential in MYCN-driven neuroblastoma^11^. Similarly, in neuroendocrine prostate cancer, EZH2 was found to act as a corepressor of MYCN target genes, and the cancer cells showed sensitivity to EZH2 methyltransferase inhibition^31^. Thus, the mechanism of cooperation between MYCN and EZH2 in these tumors seems more reliant on the canonical function of EZH2 to induce gene repression within the PRC2 complex. In sharp contrast, our data from MYCN-driven T cell lymphoma illustrates that EZH2 functions PTCL as a transcriptional activator independent from its methyltransferase function in the PRC2 complex. Furthermore, a recent study showed that EZH2 can protect MYCN from degradation by inhibiting interaction with the ubiquitin ligase FBXW7, and that this is mostly independent of its methyltransferase activity^12^. Interestingly, the authors suggest that other cancers with high expression levels of MYCN or MYC could also be dependent on high levels of EZH2^12^. Indeed, in MLL-rearranged leukemia, EZH2 was found to bind MYC via its transactivation domain, which resulted in transcriptional activation of non-PRC2 target genes, and these leukemias were strongly sensitive to EZH2 degradation^16^. In PTCL patients characterized with a high MYC signature but without overexpression of MYCN, we also observed high expression levels of EZH2 and activation of similar gene expression patterns, indicating that EZH2 could play an analogous role as transcriptional activator in this broader subgroup of PTCL.

Our findings on the oncogenic mechanisms of MYCN and EZH2 provide inspiration for improved targeted treatment strategies for PTCL patients with a high MYC expression signature. We found a striking sensitivity for HDAC inhibition in our MYCN-driven TCL mouse model, which showed synergistic effects with EZH2 degradation or dephosphorylation via CDK1 inhibition. As the FDA-approved HDAC inhibitors induce significant toxicities and are only effective in a limited group of patients, combination therapy of HDAC inhibition with EZH2-targeting compounds could improve their efficacy while decreasing their toxicity, as combination therapy will allow for lower doses.

In conclusion, we have identified MYCN as a novel oncogenic driver in PTCL, and we demonstrate its direct cooperation with EZH2 as a transcriptional co-activator of the MYCN-driven gene expression program, which can be effectively targeted by EZH2 degradation or dephosphorylation combined with HDAC inhibition.

## Acknowledgements

This study was supported by the Fund Tom Debackere for lymphoma research and by Stichting Tegen Kanker (2018/1272).

MVB and SD are supported by postdoctoral mandates for fundamental research from Stichting tegen Kanker. KD and QVT are supported by a PhD fellowship from the Research foundation Flanders (FWO). TT holds a Mandate for Fundamental and Translational Research from Stichting tegen Kanker (2019-091) and is a co-founder of the Fund ‘Me To You’ supporting research in lymphoma/leukemia. DD is supported by a postdoctoral mandate for translational research from Kom op Tegen Kanker.

We thank the KU Leuven Genomics Core, the VIB proteomics core (Sara Dufour), the VIB Histopathology Expertise Center and the VIB FACS expertise center(Jochen Lamote)for their technical services.

This study makes use of RNA-sequencing data generated by Department of Pathology and Tumor Biology, Kyoto University, as described in ref^24^.

## Authorship contributions

Conceptualization: MVB, JC, DD. Software: SD. Formal Analysis: MVB, KD, SD. Investigation: MVB, KD, SD, QVT, NieM, SP, NicM, KJ, OG, DN. Resources: SO, KK, CG, TT. Data curation: SO, TT. Writing, original draft: MVB, JC, DD. Writing, review and editing: KD, SD, QVT, NieM. Supervision: JC, DD. Funding Acquisition: MVB, DD.

