## Supplemental methods, tables and figure legend for "Aberrant MYCN expression drives oncogenic hijacking of EZH2 as a transcriptional activator in peripheral T cell lymphoma"

#### Virus production and viral transduction

Viral particles were produced by HEK293T cells using GeneJuice transfection reagent (Millipore), an ecopac packaging plasmid and the retroviral MSCV-LoxP-MYCN-LoxP-IRES-GFP expression plasmid to transfect the HEK293T cells. The supernatant carrying the viral particles was harvested and filtered after 48 hr.

For viral transduction, $1.10^6$ hematopoietic stem/progenitor cells were seeded in 1 mL RPMI 1640 supplemented with 20% fetal calf serum (Invitrogen), IL3 (10 ng/mL), IL6 (10 ng/mL), SCF (50 ng/mL) and polybrene (8 µg/mL). 1 mL viral supernatant was added, and the cells were spinfected at 2500 rpm for 90 minutes at 30°C. 4 hours after spinfection, the medium was changed into RPMI 1640 supplemented with 20% fetal calf serum (Invitrogen), IL3 (10 ng/mL), IL6 (10 ng/mL) and SCF (50 ng/mL).

#### Immunohistochemistry

Tissue samples from representative lesions were collected and fixed in 4% paraformaldehyde for 24 hours and then processed for paraffin embedding (HistoStar™ Embedding Workstation). Sections of 4 µm of thickness obtained from the paraffin-embedded tissues (Thermo Scientific Microm HM355S microtome) were mounted on Superfrost™ Plus Adhesion slides (Thermo Scientific).

Sections were routinely stained with haematoxylin and eosin (Mayers Haematoxylin, Leica, 3801582E and Eosin Y solution (aqueous), Sigma-Aldrich, HT110232-1L) for histopathological examination.

Furthermore, the PerkinElmer Opal 4-Color Manual IHC Kit (PerkinElmer/Akoya, NEL810001KT) was used for the tyramide signal amplification according to the manufacturer's protocol. For introduction of the secondary-HRP the ImmPRESS HRP Anti-Goat IgG (Peroxidase) Polymer Detection Kit (Vector Laboratories, VEC.MP-7405-15) was used for the antibody raised in goat (CD3-ε) and the OPAL Polymer HRP Ms+Rb (Akoya/Perkin Elmer, ARH1001EA) was used for the antibody raised in mouse (GFP). The various proteins were detected by using the OPAL 570 (CD3-ε) or OPAL 690 (GFP) reagents according to the manufacturer's protocol. Images were acquired on the Akoya Vectra Polaris using a x20 objective without binning. All antibodies are listed in Table S2.

### Western blotting

Cell lysates were prepared using 1X Cell Lysis Buffer (Cell Signaling Technologies) containing protease inhibitor (Complete – EDTA-free, Roche) and 1 mM $Na_3VO_4$. Proteins were separated by SDS-PAGE (NuPAGE NOVEX Bis-Tris 4–12% gels (Life Technologies) and transferred to PVDF membranes. Subsequent labelling was carried out using unlabeled primary antibodies. Western blot detection was performed with secondary antibodies conjugated with horseradish peroxidase (rabbit (GE Healthcare) or mouse (Sigma-aldrich)). Images were acquired using a cooled charge-coupled device camera system (Vilber, Fusion FX). Antibodies used for Western Blotting are listed in Table S3.

### T cell isolation

For the isolation of CD4 T cells from spleens from wild type mice, we used the MoJoSort mouse CD4 naïve T cell isolation kit (BioLegend) according to the manufacturer's instructions.

### RNA extraction

RNA from clinical samples was extracted from 4 cryosections, each 10 µm thick. The sections were resuspended in Trizol (Thermo Fisher Scientific), and after addition of chloroform, RNA was precipitated with 100% ethanol. RNA was washed and eluted with the RNeasy Mini Kit (Qiagen) according to the manufacturer's instructions.

RNA from mouse tissue was extracted using the Maxwell RSC simplyRNA Cells Kit (Promega) according to the manufacturer's instructions.

Concentrations and purity were measured with the NanoDrop 2000 (Fisher Scientific). RNA integrity was measured with the Bioanalyzer 2100 system using the RNA 6000 Nano Kit (Agilent).

### cDNA synthesis and qRT-PCR

cDNA synthesis was carried out using GoScript (Promega) and qRT-PCR was performed using the GoTaq qRT-PCR master mix (Promega) with the ViiA7 Real Time PCR system (Applied Biosystem). Primers used for qRT-PCR are listed in Table S5.

### Immunoprecipitation followed by mass spectrometry

For immunoprecipitation of EZH2, 20 µl of Magna ChIP Protein A+G Magnetic Beads (Merck Millipore) were pre-coated with 5 µL anti-EZH2 (Cell Signaling Technologies, Cat# 5246S) or rabbit IgG (Cell Signaling Technologies, Cat# 2729S) per IP, and the coated beads were mixed with pre-cleared lysates from 3 separate mice that developed MYCN-driven TCL. The beads-lysate mixtures were incubated overnight at 4 °C with gentle rotation.

Washed beads were re-suspended in 150 µl trypsin digestion buffer and incubated for 4 hours with 1 µg trypsin (Promega) at 37 ˚C. Beads were removed, another 1 µg of trypsin was added and proteins were further digested overnight at 37 ˚C. Peptides were purified on Omix C18 tips (Agilent) and dried completely in a rotary evaporator.

For LC/MS analysis, peptides were re-dissolved in 20 µl loading solvent A (0.1% trifluoroacetic acid in water/acetonitrile (ACN) (98:2, v/v)) of which 2 µl was injected for LC-MS/MS analysis on an Ultimate 3000 RSLCnano system in-line connected to a Q Exactive HF mass spectrometer (Thermo). Trapping was performed at 10 µl/min for 4 min in loading solvent A on a 20 mm trapping column (made in-house, 100 µm internal diameter (I.D.), 5 µm beads, C18 Reprosil-HD, Dr. Maisch, Germany). The peptides were separated on a 250 mm Waters nanoEase M/Z HSS T3 Column, 100Å, 1.8 µm, 75 µm inner diameter (Waters Corporation) kept at a constant temperature of 45°C. Peptides were eluted by a non-linear gradient starting at 1% MS solvent B reaching 33% MS solvent B (0.1% FA in water/acetonitrile (2:8, v/v)) in 63 min, 55% MS solvent B (0.1% FA in water/acetonitrile (2:8, v/v)) in 87 min, 99% MS solvent B in 90 minutes followed by a 10-minute wash at 99% MS solvent B and re-equilibration with MS solvent A (0.1% FA in water). The mass spectrometer was operated in data-dependent mode, automatically switching between MS and MS/MS acquisition for the 12 most abundant ion peaks per MS spectrum. Full-scan MS spectra (375-1500 m/z) were acquired at a resolution of 60,000 in the Orbitrap analyzer after accumulation to a target value of 3,000,000. The 12 most intense ions above a threshold value of 15,000 were isolated with a width of 1.5 m/z for fragmentation at a normalized collision energy of 30% after filling the trap at a target value of 100,000 for maximum 80 ms. MS/MS spectra (200-2000 m/z) were acquired at a resolution of 15,000 in the Orbitrap analyzer.

Analysis of the mass spectrometry data was performed in MaxQuant (version 2.0.1.0) with mainly default search settings including a false discovery rate set at 1% on PSM, peptide and protein level. Spectra were searched against the mouse proteins in the Reference proteins database (UP000000589, database release version of January 2021 containing 21,989 mouse protein sequences, downloaded from http://www.uniprot.org). The mass tolerance for precursor and fragment ions was set to 4.5 and 20 ppm, respectively, during the main search. Enzyme specificity was set as C-terminal to arginine and lysine, also allowing cleavage at proline bonds with a maximum of two missed cleavages. Variable modifications were set to oxidation of methionine residues, acetylation of protein N-termini. Matching between runs was enabled with a matching time window of 0.7 minutes and an alignment time window of 20 minutes. Only proteins with at least one unique or razor peptide were retained. Proteins were quantified by the MaxLFQ algorithm integrated in the MaxQuant software. A minimum ratio count of two unique or razor peptides was required for quantification. A total of 71,549 peptide-to-spectrum matches (PSMs) were performed, resulting in 10,380 identified unique peptides, corresponding to 2,105 identified proteins, of which 1,243 protein groups were reliably quantified.

### Supplemental figure legends

#### Figure S1: MYCN is aberrantly overexpressed in PTCL

(A) Normalized counts for MYCN in 481 DLBCL cases from The Cancer Genome Atlas (TCGA).

(B) Normalized counts for MYCN, MYC, MYB, MYBL1 and MYBL2 in PTCL cases from the Leuven and Kyoto cohorts.

(C) Normalized expression levels of MYCN in PTCL-NOS and in Lennert Lymphomas from the published dataset from ref[18].

(D) Volcano plot showing up- and downregulated genes in PTCL with a high MYC signature versus PTCL with a low MYC signature.

(E) Normalized counts for GATA3 and TBX21 in PTCL cases from the Leuven and Kyoto cohorts.

(F) Heatmap showing expression of genes related to the GATA3 or TBX21 subgroups of PTCL-NOS as defined by ref[17] in PTCL cases from the Leuven and Kyoto cohorts.

(G) Age at diagnosis of the PTCL patients in the Leuven cohort.

#### Figure S2: MYCN drives PTCL in a bone marrow transplant model

(A) Kaplan-Meier overall survival curve of mice transplanted with hematopoietic stem/progenitor cells (HSPCs) constitutively expressing MYCN IRES GFP.

(B) Representative FACS analysis showing the percentage of GFP-positive cells in the spleen at end-stage disease (left) and expression of myeloid, B and T cell markers (right) in mice transplanted with hematopoietic stem/progenitor cells constitutively expressing MYCN IRES GFP.

(C) Representative FACS analysis showing the percentage of GFP-positive cells and the expression of CD19 in the peripheral blood of a mouse transplanted with CD2 Cre cells transduced with the LoxP MYCN retroviral vector that developed a B cell lymphoma.

(D) Infiltration of GFP-positive cells in recipient mice transplanted with CD2 Cre cells transduced with the LoxP MYCN retroviral vector that developed B cell lymphomas.

(E) Representative FACS analysis showing the percentage of GFP-positive cells and the expression of MYCN in spleen cells of MYCN-driven T cell lymphoma.

(F) Representative FACS analysis showing the percentage of GFP-positive cells and the expression of CD3e in the spleen of recipient mouse transplanted with CD4 Cre cells transduced with the LoxP MYCN retroviral vector at end-stage disease.

(G) Representative FACS analysis showing CD8 expression from spleens of mice that developed MYCN-driven T cell lymphoma.

### Figure S3: EZH2 is a co-factor for transcriptional activation together with MYCN.

(A) Principal component analysis of RNA-sequencing data from MYCN-driven TCL and CD4 T cells.

(B) Normalized gene counts of *Cd3e*, *Dntt*, *Pdcd1*, *Cd274*, *Icos*, *Cxcr5*, *Cxcl13*, *Bcl6*, *Batf3* and *Irf4* in CD4 T cells and MYCN-driven TCL.

(C) Heatmap showing expression of T cell receptor variable genes in CD4 T cells and MYCN-driven TCL.

### Figure S4: EZH2 is a co-factor for transcriptional activation together with MYCN.

(A) Normalized expression levels of Ezh2, Ezh1, Eed and Suz12 in CD4 T cells and in MYCN-driven TCL cells.

(B) Representative ChIP-seq tracks for Ezh1, Suz12 and Eed showing binding of MYCN, Ezh2, Suz12, p300 and histone marks H3K4me3, H3K27ac and H3K27me3 in MYCN-driven TCL cells.

(C) Venn diagram of EZH2 peaks in direct MYCN target genes and their overlap with H3K27ac or H3K27me3 peaks.

(D) Representative ChIP-seq tracks of the Aurora Kinase A (Aurka) promoter region showing binding of MYCN, Ezh2, Suz12, p300 and histone marks H3K4me3, H3K27ac and H3K27me3 in MYCN-driven TCL. The 1513 bp region cloned into the pGL4.27 luciferase plasmid is indicated within the black bar.

(E) Luciferase assay in HEK293T cells showing transcriptional activation of Firefly luciferase by MYCN and/or EZH2, using the pGL4.27 luciferase plasmid with and without the Aurora Kinase A promoter region.

(F) Luciferase assay in HEK293T cells showing transcriptional activation of Firefly luciferase by MYCN and/or EZH2 (WT or Y244D mutant).

(G) Dose response curve for the EZH2 methyltransferase inhibitor Tazemetostat on *ex vivo* cultured MYCN-driven TCL cells.

(H) Dose response curve for the EZH2 methyltransferase inhibitor Valemetostat on *ex vivo* cultured MYCN-driven TCL cells.

(I) Mass spectrometry spectrum of the identified Ezh2 phospho-site on Threonine 487.

### Figure S5: MYCN-driven TCL is sensitive to EZH2 depletion and HDAC inhibition.

(A) MYCN-driven T cell lymphoma cells, non-malignant CD4 T cells or TAL1–AKT leukemia/lymphoma cells treated *ex vivo* with DMSO or 500 nM romidepsin for 4h.

(B) Dose response curve for the EZH2 degrader MS1943 with or without simultaneous JQ1 treatment on *ex vivo* cultured MYCN-driven TCL cells.

(C) Synergy matrix plot showing δ-scores for *ex vivo* cultured MYCN-driven TCL cells treated with MS1943 + JQ1 (max ZIP synergy score = maximal score for a specific dose combination; average ZIP synergy score = the average δ-score for the whole range of concentrations shown in the synergy matrix).

(D) Dose response curve for the CDK1 inhibitor Ro-3306 with or without simultaneous JQ1 treatment on *ex vivo* cultured MYCN-driven TCL cells.

(E) Synergy matrix plot showing δ-scores for *ex vivo* cultured MYCN-driven TCL cells treated with Ro-3306 + JQ1 (max ZIP synergy score = maximal score for a specific dose combination; average ZIP synergy score = the average δ-score for the whole range of concentrations shown in the synergy matrix).

### Supplemental Tables

*Table S1: antibodies used for Flow cytometry*

| Antibody | Fluorophore | Supplier | Cat# |
|---|---|---|---|
| Armenian Hamster monoclonal anti-CD3ε [clone 145-2C11] | Pacific Blue | Biolegend | 100334 |
| Rat monoclonal anti-CD4 [clone GK1.5] | PE-Vio770 | Miltenyi Biotec | 130-124-712 |
| Rat monoclonal anti-CD8a | VioBlue | Miltenyi Biotec | 130-102-431 |
| Rat monoclonal anti-CD4 | VioBlue | Miltenyi Biotec | 130-101-962 |
| Rat monocloncal anti-CD19 [clone eBio1D3] | PE-Cy7 | Thermo Fischer Scientific | 25-0193-81 |
| Rat monoclonal Anti-Cd11b | APC-vio770 | Miltenyi Biotec | 130-096-834 |
| Rat monoclonal anti-B220 | APC-vio770 | Miltenyi Biotec | 130-102-267 |
| Rat monoclonal anti-B220 | VioBlue | Miltenyi Biotec | 130-118-321 |
| Anti-phospho-EZH2 (Thr487) | / | Invitrogen | PA5-105660 |
| Rabbit Anti-IgG | Alexa Fluor 405 | Abcam | ab175652 |

*Table S2: antibodies used for immunohistochemistry*

| Antibody | Supplier | Cat# |
|---|---|---|
| Goat polyclonal anti-CD3e | Santa Cruz | sc-1127 |
| Mouse monoclonal anti-GFP | Santa Cruz | sc-9996 |

*Table S3: antibodies used for Western blotting*

| Antibody | Supplier | Cat# |
|---|---|---|
| Mouse polyclonal anti-MYCN | Cell Signaling Technologies | 9405S |
| Rabbit monoclonal anti-EZH2 [clone D2C9] | Cell Signaling Technologies | 5246S |
| Rabbit monoclonal anti-SUZ12 [clone D39F6] | Cell Signaling Technologies | 3737S |
| Mouse monoclonal Anti-β–Actin [Clone AC-15] | Merck | A5441 |

*Table S4: antibodies used for ChIP-sequencing*

| Antibody | Supplier | Cat# |
|---|---|---|
| Mouse polyclonal anti-MYCN | Cell Signaling Technologies | 9405S |
| Rabbit monoclonal anti-EZH2 [clone D2C9] | Cell Signaling Technologies | 5246S |
| Rabbit monoclonal anti-SUZ12 [clone D39F6] | Cell Signaling Technologies | 3737S |
| Rabbit monoclonal anti-p300 [clone D2X6N] | Cell Signaling Technologies | 54062S |
| Rabbit polyclonal anti-H3K27ac | Abcam | ab4729 |
| Rabbit monoclonal anti-H3K27me3 [clone C36B11] | Cell Signaling Technologies | 9733S |
| Rabbit monoclonal anti-H3K4me3 [clone C42D8] | Cell Signaling Technologies | 9751S |

*Table S5: qRT-PCR primers*

| Target | Primer direction | Sequence |
|---|---|---|
| Mouse Ezh2 | Fw | CTGTCTCACGTGTGGAGCTG |
| | Rv | AGACGGTGCCAGCAGTAAGT |
| Mouse Myb | Fw | GCTGGAGTTGCTCCTGATGT |
| | Rv | GCTGCAAGTGTGGTTCTGTG |
| Mouse Foxm1 | Fw | TGGAGCAGAATCGGGTTAAG |
| | Rv | GTGCTGTTGATGGCAAACTG |
| Mouse Cdk6 | Fw | GACCTCTGGAGTGTCGGTT |
| | Rv | GTCCAATGATGTCCAAGATTTTTCC |

*Table S6: compounds used for ex vivo treatments*

| Compound | Supplier | Cat# | Cas# |
|---|---|---|---|
| MS1943 | MedKoo | 462549 | 2225938-17-8 |
| Romidepsin | AdooQ Bioscience | A11920 | 128517-07-7 |
| Tazemetostat | Selleckchem | S7128 | 1403254-99-8 |
| Valemetostat | Selleckchem | S8926 | 1809336-39-7 |
| Ro-3306 | Medchem Express | HY-12529 | 872573-93-8 |
| Belinostat | Selleckchem | S1085 | 414864-00-9 |
| JQ1 | Selleckchem | S7110 | 1268524-70-4 |
