## Supplementary figures and images for "Aberrant MYCN expression drives oncogenic hijacking of EZH2 as a transcriptional activator in peripheral T cell lymphoma"

### Figure S1

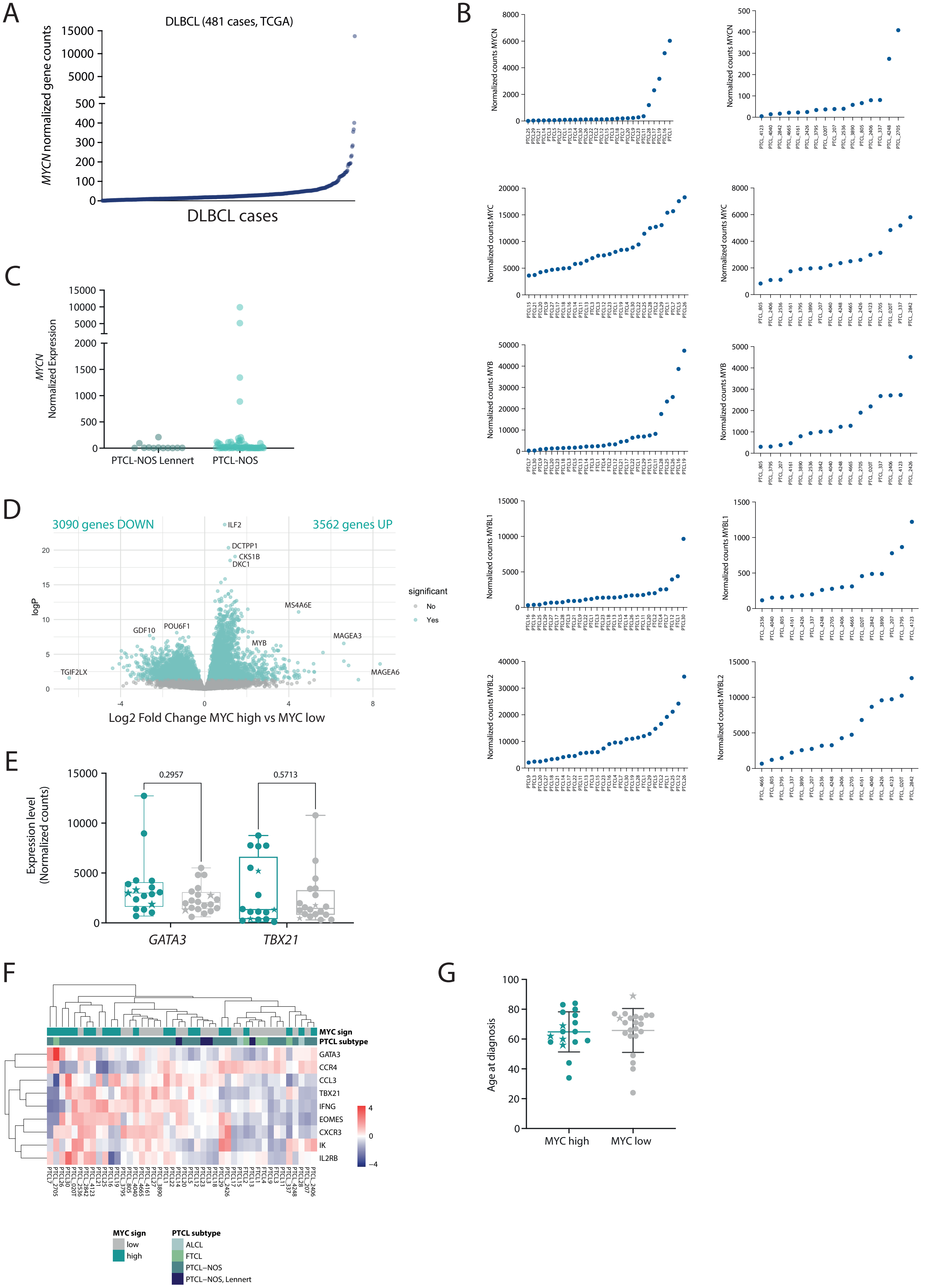

### Figure S2

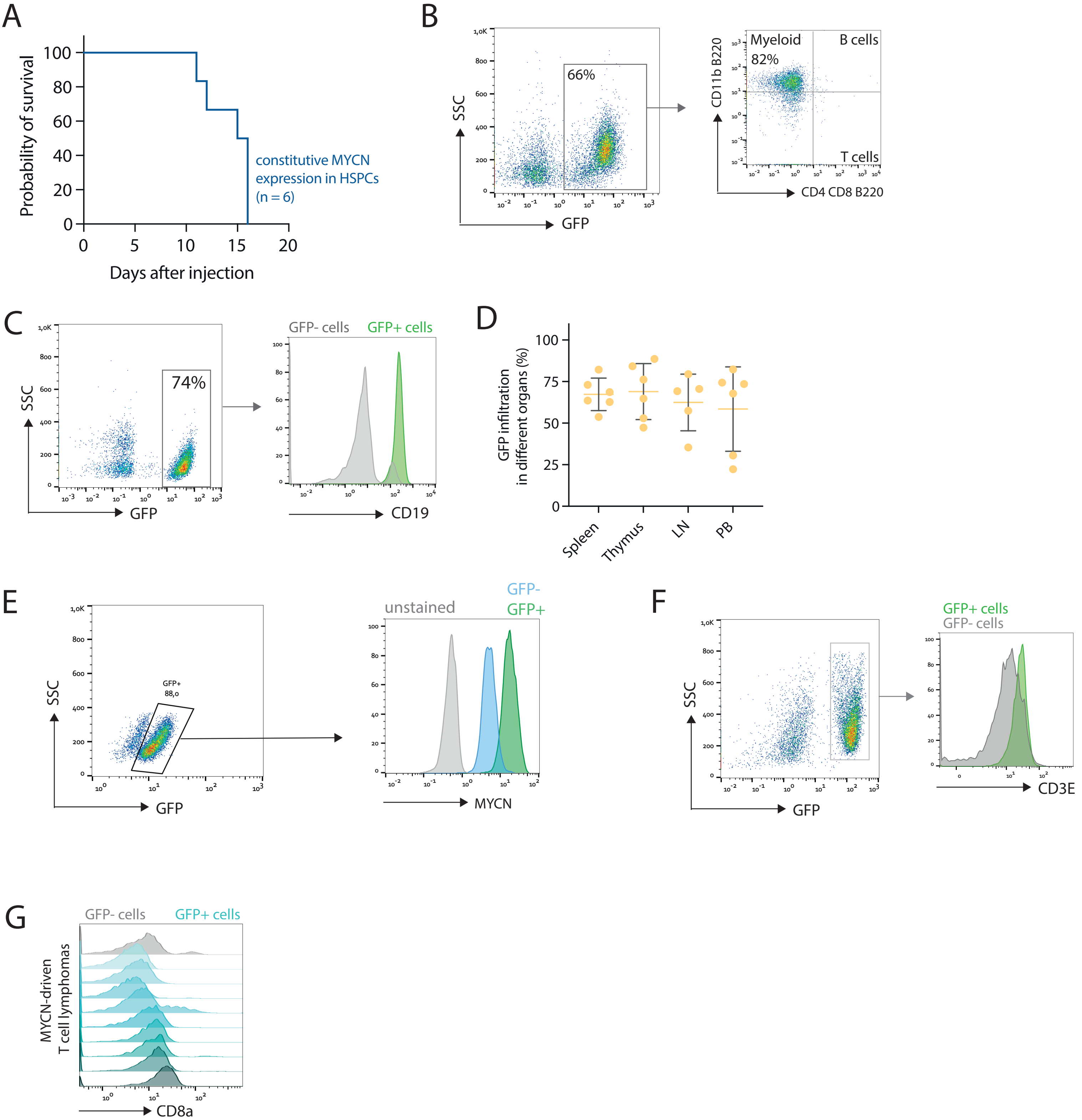

### Figure S3

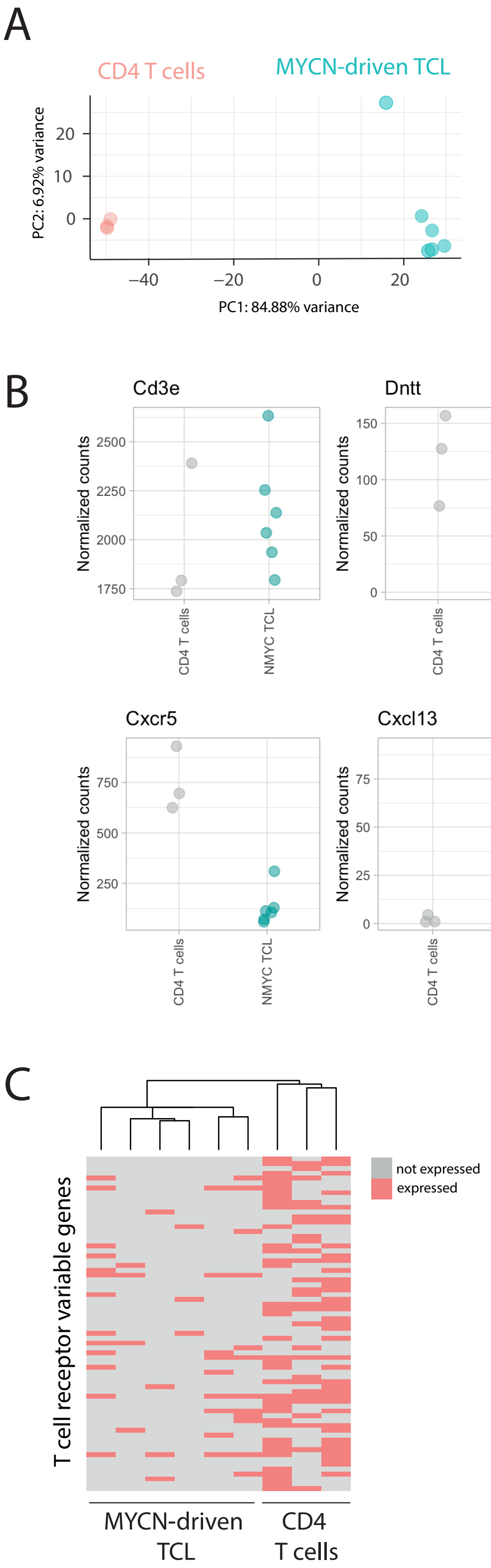

### Figure S4

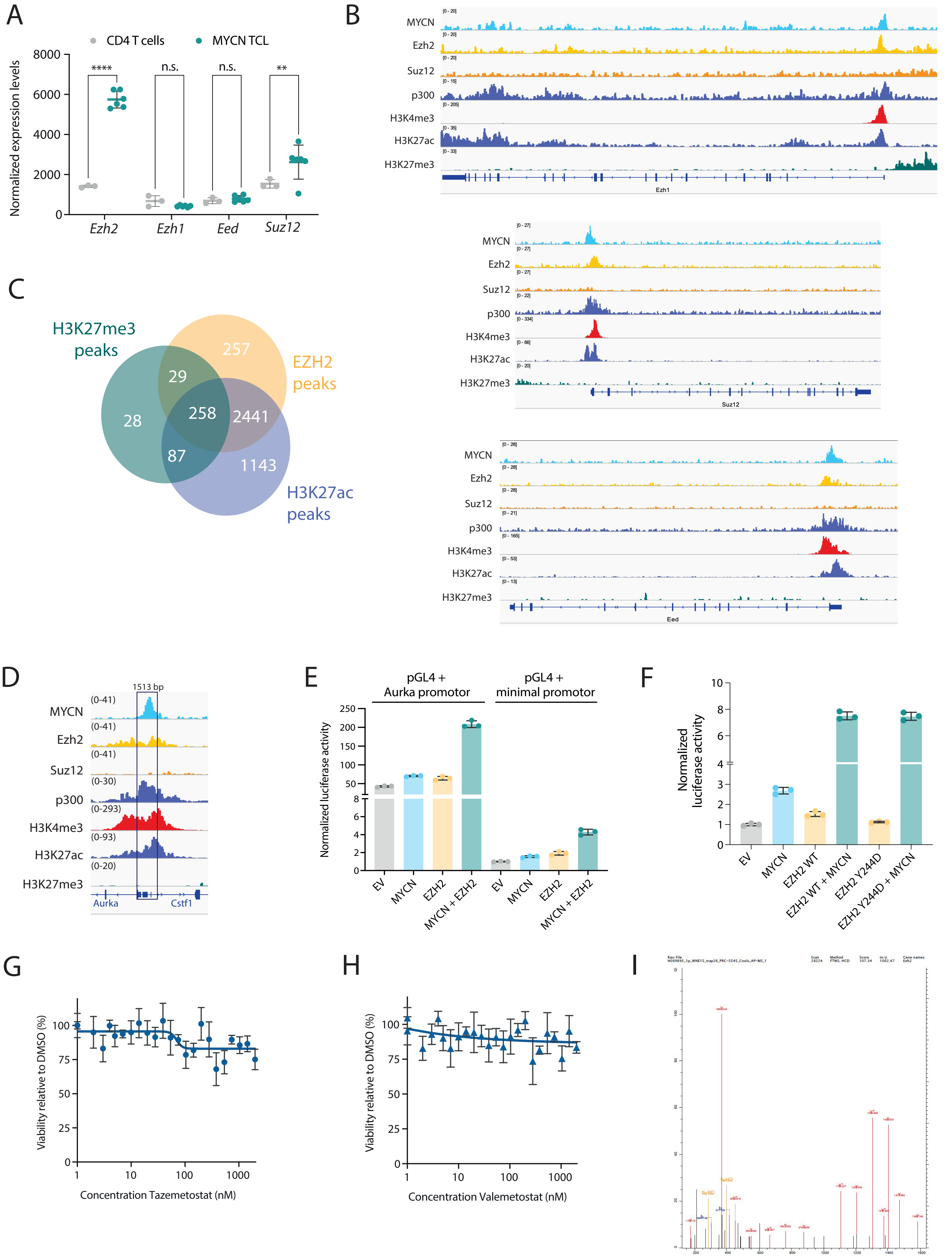

### Figure S5

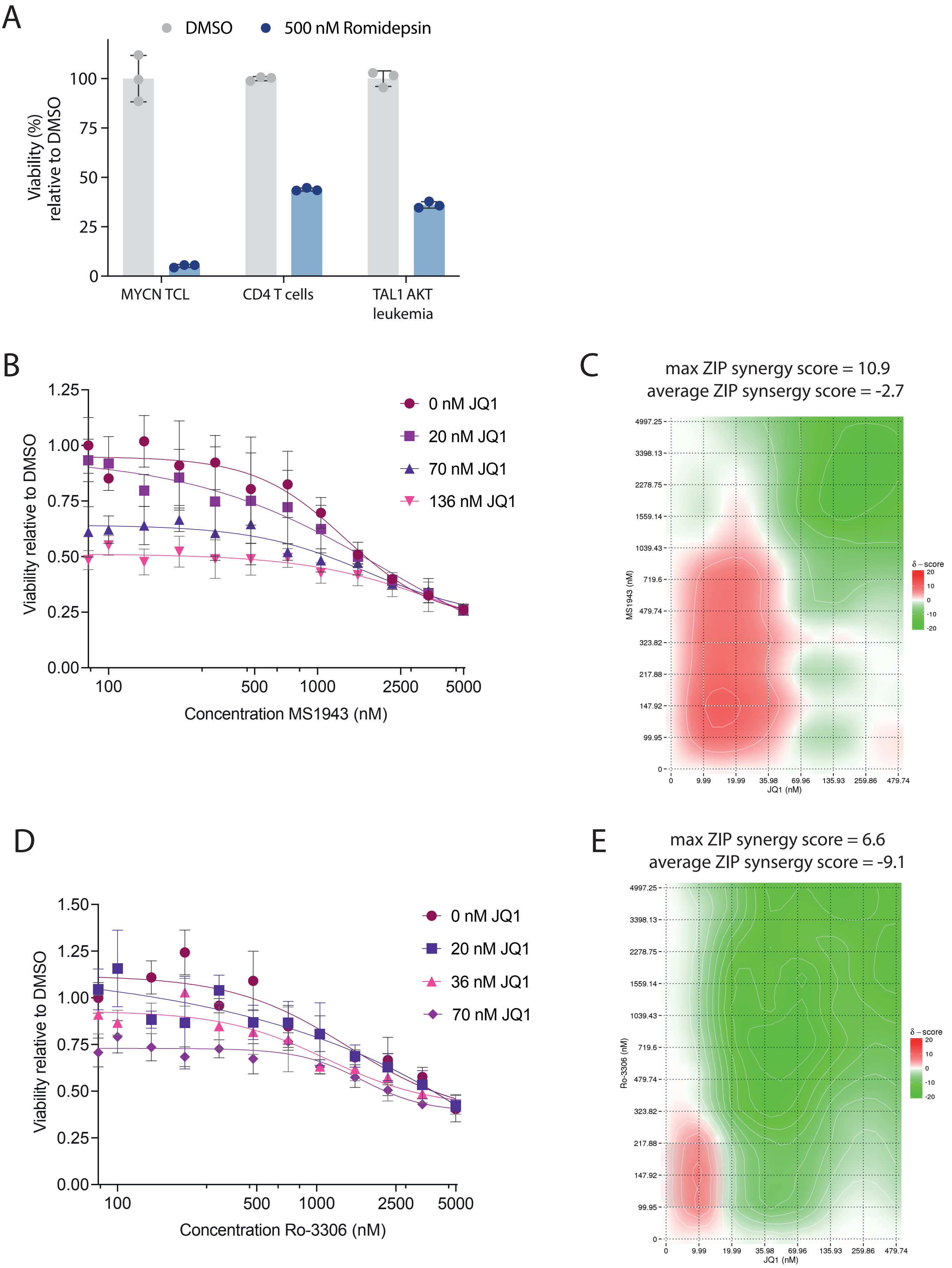
